# Inducible expression of human *C9ORF72* 36x G_4_C_2_ hexanucleotide repeats is sufficient to cause RAN translation and rapid muscular atrophy in mice

**DOI:** 10.1101/2020.09.15.297259

**Authors:** F.W. Riemslagh, E.C. van der Toorn, R.F.M Verhagen, A. Maas, L.W.J. Bosman, R.K. Hukema, R. Willemsen

## Abstract

The hexanucleotide G_4_C_2_ repeat expansion in the first intron of the *C9ORF72* gene explains the majority of frontotemporal dementia (FTD) and amyotrophic lateral sclerosis (ALS) cases. Numerous studies have indicated the toxicity of dipeptide repeats (DPRs) which are produced via repeat-associated non-AUG (RAN) translation from the repeat expansion and accumulate in the brain of C9FTD/ALS patients. Mouse models expressing the human *C9ORF72* repeat and/or DPRs show variable pathological, functional and behavioral characteristics of FTD and ALS. Here, we report a new Tet-on inducible mouse model that expresses 36x pure G_4_C_2_ repeats with 100bp upstream and downstream human flanking regions. Brain specific expression causes the formation of sporadic sense DPRs aggregates upon 6 months dox induction but no apparent neurodegeneration. Expression in the rest of the body evokes abundant sense DPRs in multiple organs, leading to weight loss, neuromuscular junction disruption, myopathy and a locomotor phenotype within the time frame of four weeks. We did not observe any RNA foci or pTDP-43 pathology. Accumulation of DPRs and the myopathy phenotype could be prevented when 36x G_4_C_2_ repeat expression was stopped after 1 week. After 2 weeks of expression, the phenotype could not be reversed, even though DPR levels were reduced. In conclusion, expression of 36x pure G_4_C_2_ repeats including 100bp human flanking regions is sufficient for RAN translation of sense DPRs and evokes a functional locomotor phenotype. Our inducible mouse model highlights the importance of early diagnosis and treatment for C9FTD/ALS patients.

**Summary statement:** Only 36 C9ORF72 repeats are sufficient for RAN translation in a new mouse model for ALS and FTD. Reducing toxic dipeptides can prevent but not reverse the phenotype.

## Introduction

FTD is a neurological disease characterized by neuronal loss in the frontal and temporal lobes leading to behavioral and personality changes and language deficits(Hernandez et al., 2018; Woollacott and Rohrer, 2016). The prevalence of FTD is approximately 15-20 cases per 100.000 people and the age of onset is usually between 45 to 65 years(Woollacott and Rohrer, 2016). FTD is part of a disease spectrum that also comprises ALS(Couratier et al., 2017; Strong et al., 2017). ALS is a rapid progressive motor neuron disorder that affects the upper motor neurons in the motor cortex and the lower motor neurons in the anterior horn of the spinal cord(Grad et al., 2017; Oskarsson et al., 2018). ALS patients develop muscle weakness, spasticity, atrophy and eventually paralysis(Grad et al., 2017; Oskarsson et al., 2018). The prevalence of ALS is about 5 in 100.000 people and the age of onset is between 50 and 60 years of age(Grad et al., 2017; Oskarsson et al., 2018). The hexanucleotide G_4_C_2_ repeat expansion in the *C9ORF72* gene explains almost 90% of the families presenting with both FTD and ALS symptoms(DeJesus-Hernandez et al., 2011; Renton et al., 2011) (referred to as C9FTD/ALS). Patients can be mosaic for repeat size and often have longer repeats in brain tissue than in DNA isolated from blood samples (Fournier et al., 2019; Nordin et al., 2015; van Blitterswijk et al., 2013). So far, repeat sizes of 24 – 4400 have been reported(Iacoangeli et al., 2019; Van Mossevelde et al., 2017). Associations between repeat size and clinical diagnosis have not resulted in a clear picture (Fournier et al., 2019; Gijselinck et al., 2016; van Blitterswijk et al., 2013). Thus, the exact repeat size that triggers disease onset is not known.

Three mechanisms for the *C9ORF72* repeat expansion have been proposed to cause C9FTD/ALS(Balendra and Isaacs, 2018): 1) Hypermethylation of the repeat and surrounding CpG islands can lead to reduced levels of the normal C9ORF72 protein(Belzil et al., 2013; Waite et al., 2014). *C9orf72* knock-out mice have shown its essential function in immunity, but do not present with FTD or ALS symptoms(Balendra and Isaacs, 2018). However, haploinsufficiency can still modify the effects of gain-of-function mechanisms via the normal cellular function of the C9ORF72 protein in autophagy and lysosomal biogenesis(Sellier et al., 2016; Shi et al., 2018). 2) Repeat containing RNA from both sense and antisense direction can form secondary structures(Kovanda et al., 2015; Su et al., 2014) and RNA foci(Gendron et al., 2013; Mizielinska et al., 2013). Repeat–containing RNA or RNA foci can sequester RNA-binding proteins and prevent their normal functioning in the cell(Haeusler et al., 2016). 3) The G_4_C_2_ repeat can also be translated into dipeptide repeats (DPRs) via repeat-associated non-ATG (RAN) translation (Ash et al., 2013; Gendron et al., 2013; Mori et al., 2013). RAN translation occurs in all reading frames of sense and antisense transcripts and results in the formation of poly-glycine-alanine (GA), poly-glycine-proline (GP), poly-glycine-arginine (GR), poly-proline-alanine (PA) and poly-proline-arginine (PR). DPRs have been found throughout the brains of C9FTD/ALS patients(Mackenzie et al., 2015) and especially poly-GR has been associated with neurodegeneration(Gittings et al., 2020; Saberi et al., 2018; Sakae et al., 2018). Multiple cell and animal models have indicated the detrimental effect of expression of both arginine-containing DPRs poly-GR and poly-PR and the slightly less toxic poly-GA(Balendra and Isaacs, 2018; Boeynaems et al., 2016; Jovicic et al., 2015; Kanekura et al., 2016; Kwon et al., 2014; Mizielinska et al., 2014; Swaminathan et al., 2018; Tao et al., 2015; Wen et al., 2014; Yamakawa et al., 2015; Yang et al., 2015). So far, 11 loss-of-function and 10 gain-of-function *C9ORF72* mouse models have been published (reviewed in (Balendra and Isaacs, 2018; Batra and Lee, 2017)), and 5 DPR-only mouse models investigating the role of poly-GA(Schludi et al., 2017; Zhang et al., 2016), poly-GR(Zhang et al., 2018) and poly-PR(Hao et al., 2019; Zhang et al., 2019). All mouse models support a gain-of-function hypothesis in C9FTD/ALS, although not all BAC mice show neurodegeneration or a motor phenotype associated with ALS (reviewed in (Balendra and Isaacs, 2018; Batra and Lee, 2017)). The effect of DPRs has been studied extensively (reviewed in(Balendra and Isaacs, 2018)), but possible reversibility and the exact number of repeats needed for RAN translation *in vivo* are still not determined(Cleary et al., 2018; Green et al., 2016).

Here, we describe a new mouse model that expresses human *C9ORF72* 36x pure G_4_C_2_ repeats with 100bp upstream and downstream human flanking regions under the expression of an inducible Tet-on promotor. This system allows for temporal and spatial expression of the repeat expansion. Expression of 36x pure G_4_C_2_ repeats was sufficient to produce sense DPRs and a locomotor phenotype upon four weeks of induction of expression. In order to study possible reversibility, expression was stopped after 1 or 2 weeks followed by a washout period of 2-3 weeks to prevent further build-up and subsequent reduction of the amount of DPRs.

## Results

### Generation and expression pattern of the human 36x G_4_C_2_ repeat mouse model

We generated our mouse model from DNA isolated from a C9FTD patient’s blood and amplified the repeat in three consecutive PCR rounds using primers that flanked the *C9ORF72* repeat expansion (for primer sequences see materials and methods). The PCR product was cloned into TOPO and subsequently into a Tet-on vector with a GFP reporter gene(Hukema et al., 2014) (figure 1A). Sequencing of this construct revealed a repeat size of 36x pure G_4_C_2_ repeats with 118 bp upstream and 115 bp downstream human flanking region (supplementary figure 1). The transgene (containing the TRE promotor, 36x G_4_C_2_ repeats and the GFP gene) was injected into pronuclei of C57BL/6J mice. Founder mice were screened for the presence and size of the transgene and transmission to their offspring. Genotyping for transgene presence was performed with primers located 5’ of the repeat. Repeat size estimation was established using the Asuragen *C9ORF72* repeat kit that is also used in routine human diagnostics. Repeat size remained stable between generations and between multiple organs of the same mouse (figure 1B). Transgenic mice were born at Mendelian frequencies and showed normal viability.

**Figure 1:**
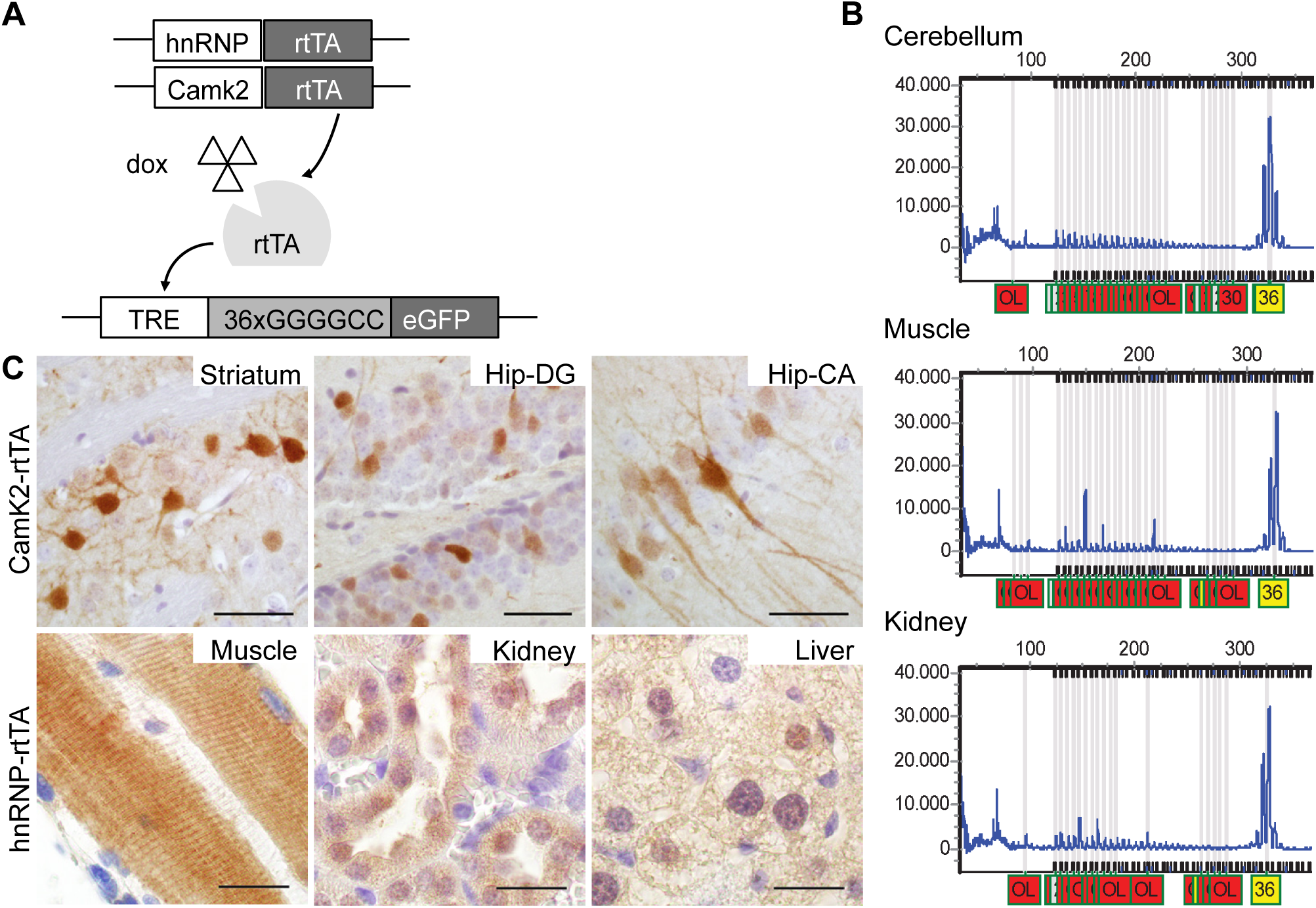
Generation and expression of the 36x G_4_C_2_ repeat mouse model. A) Schematic of the Tet-on system. Mice either have a Camk2-alpha-rtTA or hnRNP-rtTA transgene that expresses rtTA in a brain specific manner or in the whole body. Upon binding of doxycycline, rtTA can bind the TRE response element and start transcription of the C9ORF72 G4C2 repeat expansion and GFP gene, which has its own start site. During the whole study, single transgenic littermates were used as control and received the same dox treatment. B) DNA isolated from different tissues from the same mouse (17129-5 ladder mouse 4 weeks on dox) and analyzed with the Asuragen C9ORF72 PCR kit shows a repeat length of 36 in all tissues. This was done for at least 3 independent mice. C) Upper panel: GFP expression was detected in striatum and hippocampus cornu ammonis and hippocampus dentate gyrus in double transgenic Camk2-alpha-rt-TA/TRE-36G4C2-GFP mice after dox administration. Lower panel; GFP expression in EDL muscle, kidney and liver of double transgenic hnRNP-rtTA/TRE-36xG4C2-GFP mice. GFP staining was performed on all mice in this study. ST 4 weeks dox n=15, DT 4 weeks dox n=16. Scale bars are 20 µm.

Heterozygous transgenic mice containing the TRE-36xG_4_C_2_–GFP construct were bred with two different heterozygous rtTA driver lines. We choose the CamK2α (Ca^2+^/calmodulin-dependent protein kinase II linked reverse tetracycline-controlled trans-activator) rtTA driver because of its validated expression in the hippocampus and cortex, brain areas that are showing pathology in C9FTD/ALS patients(Odeh et al., 2011). To study expression in the rest of the body we used a hnRNP (heterogeneous nuclear ribonucleoprotein 2B1) rtTA driver. The resulting litters consist of 4 different genotypes referred to in the rest of the paper as double transgenic (containing both the TRE-36xG_4_C_2_–GFP construct and one of the rtTA constructs) or single transgenic littermates (containing only the TRE or only an rtTA-driver construct). Wildtype littermates were not used in this study. Mice were administered doxycycline (dox) in their drinking water at 6 weeks of age to turn on transgene expression, which revealed specific expression of GFP in the double transgenic (DT) mice only and no transgene expression in single transgenic (ST) mice (supplementary figure 2). Using the hnRNP-rtTA driver, we observed expression in almost all tissues, including extensor digitorum longus (EDL) muscle, liver, kidney (figure 1C), heart and lung (supplementary figure 2), but not in brain and spinal cord after max. 4 weeks of dox treatment (supplementary figure 2). Using the Camk2-alpha-rtTA driver, we observed GFP expression only in striatum and hippocampus dentate gyrus and cornu ammonis, as expected for this driver (figure 1C). GFP expression was detectable after 1 week of dox administration and remained detectable over 6 months (data not shown).

**Figure 2:**
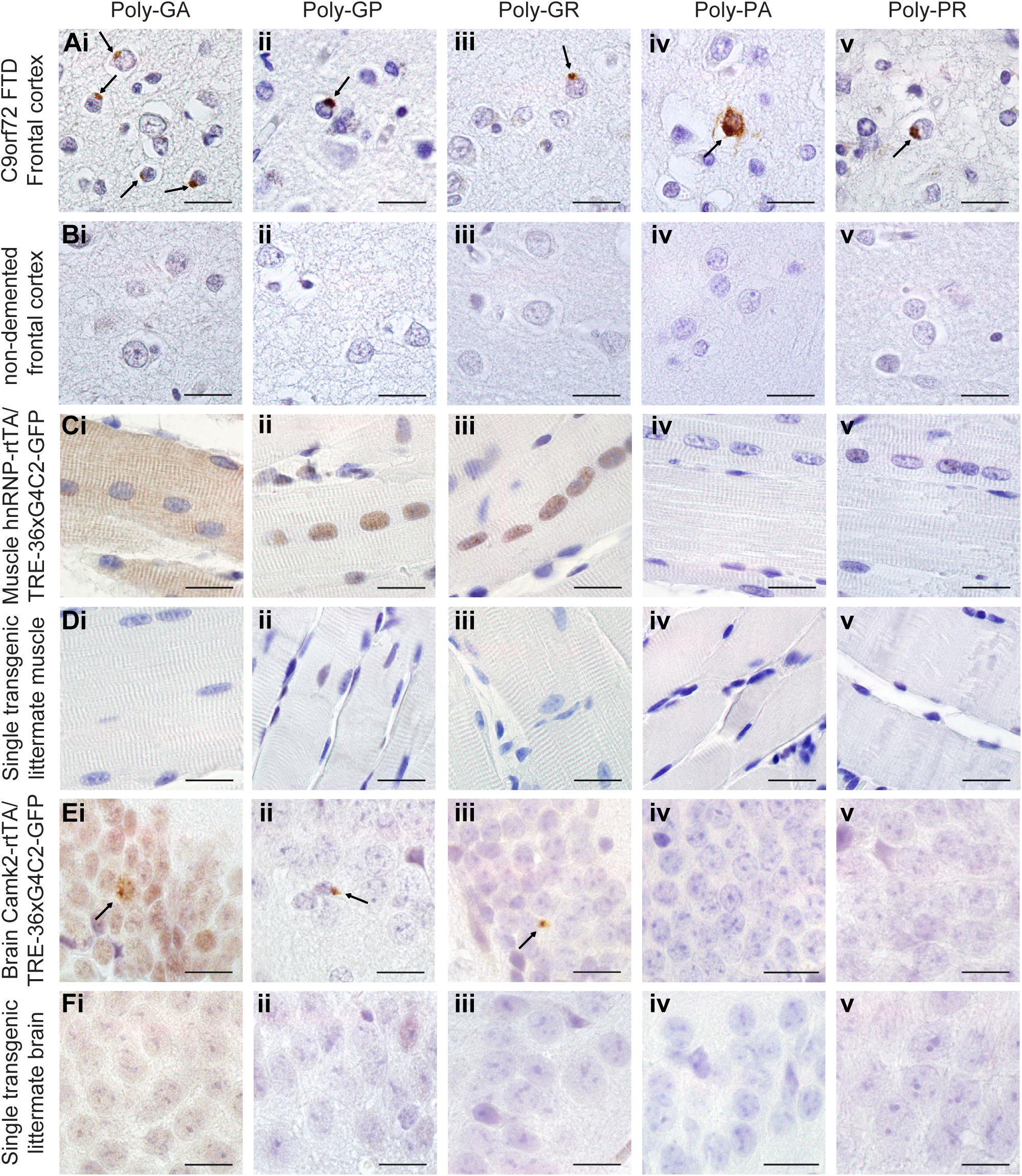
Expression of 36x G_4_C_2_ human repeats is enough to evoke sense DPR formation *in vivo*. Human prefrontal cortex of A) C9FTD patients or B) non-demented controls were used as positive and negative control for detection of DPR pathology. C) In TRE-36xG4C2-GFP/hnRNP-rtTA double transgenic mice, poly-GA shows both diffuse cytoplasmic and nuclear localization, while diffuse poly-GP and poly-GR are observed in the nucleus of the EDL muscle. E)TRE-36xG4C2-GFP/Camk2-alpha-rtTA double transgenic mice show some sparse perinuclear aggregates of sense DPRs in the hippocampus dentate gyrus (pointed at by arrows). D) and F) Single transgenic littermates received the same dox treatment and contained only one transgene either TRE-only or rtTA-only, are all negative for DPRs. DPR staining was performed on all mice in this study. ST 4 weeks dox n=15, DT 4 weeks dox n=16. All scale bars are 20 µm.

### Human 36x G_4_C_2_ repeat mice show DPR expression but no RNA foci

To further characterize the expression of the transgene in our mouse model, we performed fluorescence in situ hybridization (FISH) to test for the presence of sense and antisense RNA foci. Unfortunately, we were unable to detect RNA foci in multiple organs at 1-24 weeks of age in any of the driver lines (supplementary figure 3), despite the fact that our protocol was optimized to detect RNA foci in post-mortem human C9FTD/ALS frontal cortex paraffin tissue used as positive control (supplementary figure 3). Sense transcribed DPRs (poly-GA, -GP and -GR) were present in all GFP-positive tissues of DT hnRNP-rtTA mice (figure 2). We did not observe antisense DPRs in mice (figure 2 C-F), although we could detect them in C9FTD/ALS patient frontal cortex sections that were used as positive control (figure 2 A-B). To validate that no antisense transcripts were produced in our mouse model, we performed rt-PCR (supplementary figure 4). Again, antisense transcripts were only observed in RNA isolated from frozen frontal cortex of a human C9FTD patient (supplementary figure 4).

**Figure 3:**
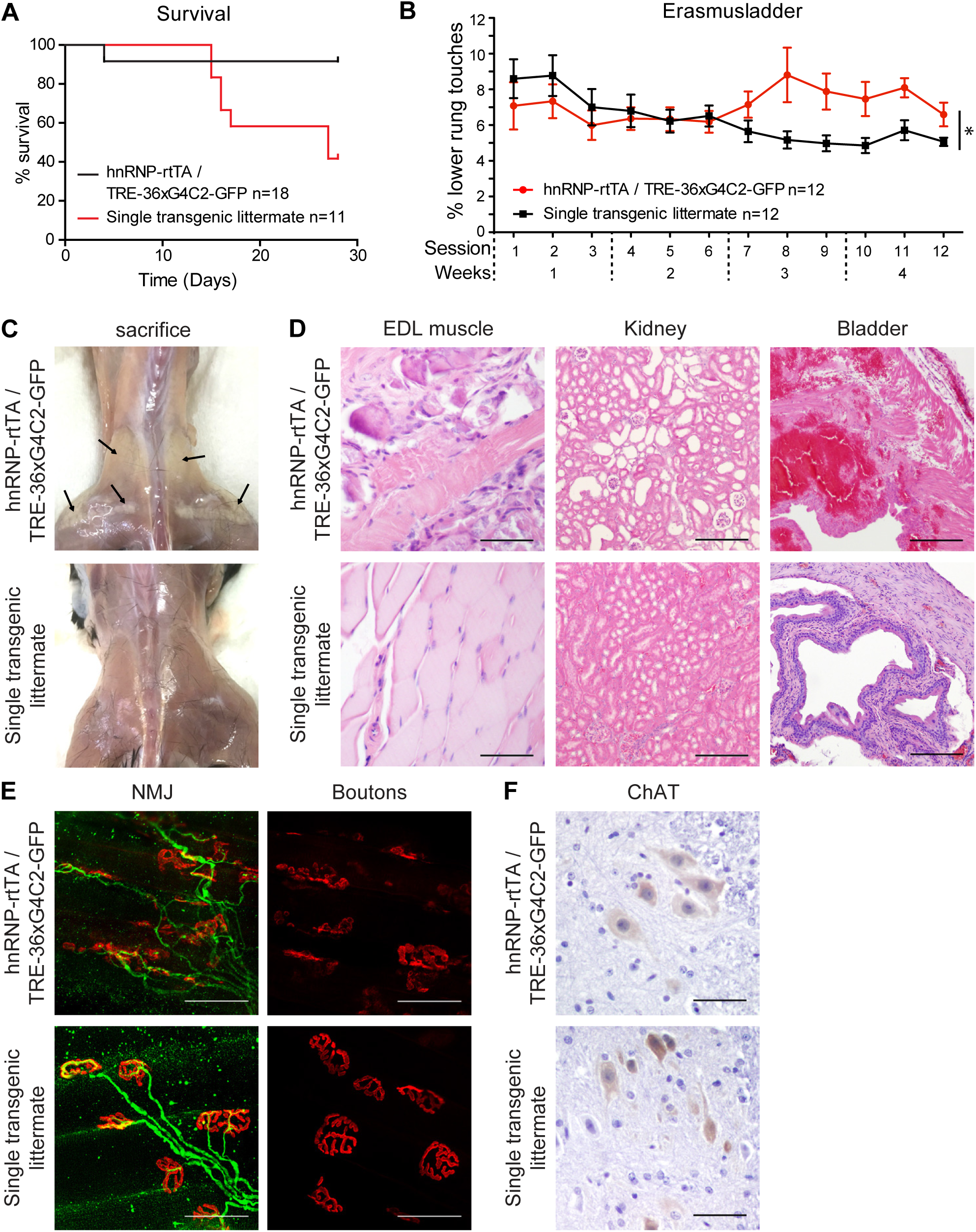
Expression of 36x G_4_C_2_ human repeats *in vivo* causes a locomotor phenotype and muscular dystrophy within 4 weeks. A) TRE-36xG4C2-GFP/hnRNP-rtTA double transgenic mice that receive dox show reduced survival after 1-3 weeks (n=18) compared to single transgenic littermates (containing only one transgene either TRE-only or rtTA-only) who received the same dox treatment (n=11). B) Mice that survive develop a locomotor phenotype on the Erasmusladder after 7 sessions (3 sessions/week). N = 12 mice per group. Note that from session 7 on, some mice had to be excluded due to severe pathology or incapability to cope with the behavioral assay. Session 12 includes data from N = 10 single transgenic control and N = 8 double transgenic mice. All mice received the same dox treatment. Two-way ANOVA p=0.0001 for genotype and p<0.0001 for the interaction between genotype and time. Error bars represent standard error of the mean (SEM) C) Sacrificed DT mice show white appearance of back and upper leg muscles. D) HE staining of the EDL muscle, kidney and bladder of DT mice. Scale bars of EDL and kidney images are 50 µm. Scale bar of bladder image is 200 µm. E) NMJ staining of the EDL muscle shows dissolving boutons (red, α-bungarotoxin) and disorganized axonal projections (green, neurofilament antibody). Scale bar 50 µm. F) The number of ChAT-positive motor neurons in the spinal cord does not differ between DT and ST control littermates. Scale bar is 20 µm. All stainings were performed on all mice in this study. ST 4 weeks dox n=15, DT 4 weeks dox n=16.

**Figure 4:**
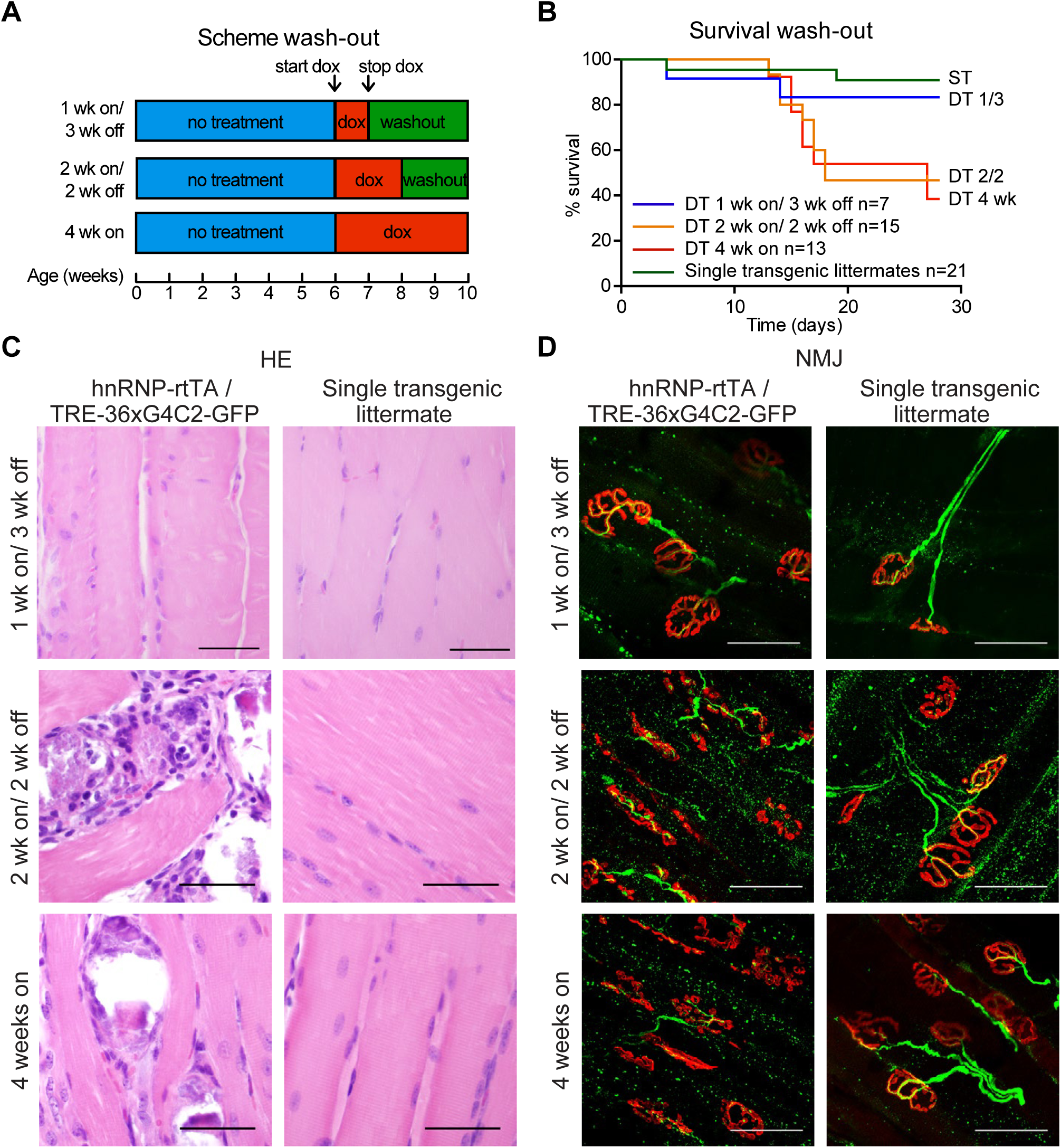
Early dox withdrawal can prevent but not reverse the muscular dystrophy and NMJ phenotype of TRE-36xG4C2-GFP/hnRNP-rtTA double transgenic mice. A) Scheme of different washout groups. B) Survival curve for washout experiment. Mice that received 1 week dox followed by 3 weeks washout (1 wk on/ 3 wk off n=7) show high survival. Two weeks dox followed by 2 weeks washout (2 wk on/ 2wk off n=15) shows the same reduction in survival as 4 weeks continuous dox administration (4 wk on n=13). Single transgenic littermates (containing only one transgene either TRE-only or rtTA-only), were distributed over the groups and received the same dox treatment (n=21). C) HE staining of the EDL muscle remains normal in the 1 wk on/ 3 wk off group but shows distortion in TRE-36xG4C2-GFP/hnRNP-rtTA DT mice that received 2 weeks dox followed by 2 weeks of normal drinking water. Scale bars are 50 µm. D) NMJ of the EDL muscle shows collapsed boutons (red, α-bungarotoxin) and axonal projections (green, neurofilament antibody) in DT groups that receive 2 or more weeks dox. Scale bars 50 µm. All stainings were performed on all mice in this study. Numbers per group are: ST 1 week dox n=7, DT 1 week dox n=8, ST 1 week on/3 weeks off n=4, DT 1 week on/3 weeks off n=7, ST 2 weeks dox n=6, DT 2 weeks dox n=8, ST 2 weeks on/2 weeks off n=6, DT 2 weeks on/2 weeks off n=5, ST 4 weeks dox n=15, DT 4 weeks dox n=16

Interestingly, sense DPRs differed in subcellular localization. Poly-GA was visible as diffuse nuclear and cytoplasmic labeling in DT hnRNP-rtTA mice, while poly-GP and –GR were only observed in the nucleus (figure 2C-D and supplementary figure 5). In the DT Camk2-alpha-rtTA mice, we could detect some small perinuclear aggregates in the striatum and hippocampus after 24 weeks of dox administration (figure 2E-F). However, the numbers of aggregates were very rare (about 1 aggregate per sagittal brain section). Longer follow-up of these mice is not possible, as administration of dox for more than 6 months often leads to intestine problems and unacceptable welfare discomfort for the mice. Both Camk2- and hnRNP-driven 36xG_4_C_2_ repeat mice show no abundant pathological hallmarks of C9FTD/ALS, including p62 and pTDP-43 aggregates in brain and muscle (supplementary figure 6). Nor did we observe any signs of neurodegeneration (cleaved caspase-3 staining, supplementary figure 6), astrogliosis or microgliosis (supplementary figure 7). As DPR inclusions were very rare in Camk2-alpha-rtTA mice and expression of DPRs was evident in the hnRNP-rtTA mice, we choose to focus on the DT hnRNP-rtTA mice for further assessment of the toxic effect of DPR expression in multiple organs *in vivo*.

**Figure 5:**
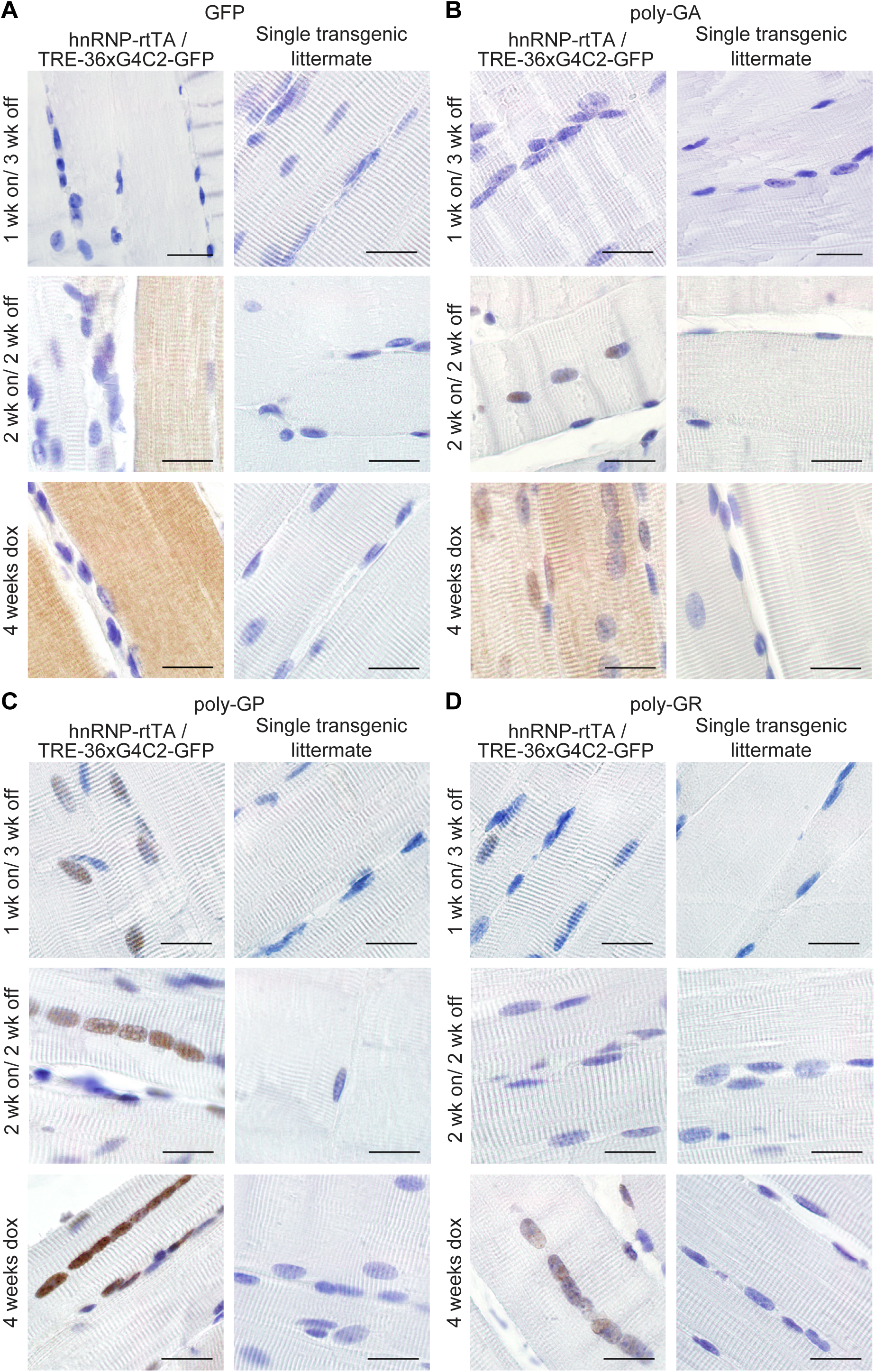
GFP and sense DPRs are reduced after dox withdrawal. A) GFP staining on EDL muscle of TRE-36xG4C2-GFP/hnRNP-rtTA double transgenic mice shows reduction in the intensity of staining when mice received 1 or 2 weeks of dox water followed by 2-3 weeks of normal drinking water compared to DT littermates that received 4 weeks of dox. B) Poly-GA staining of EDL muscle shows clearance of cytoplasmic poly-GA in all washout groups but retention of nuclear poly-GA after two weeks of dox withdrawal. C) Poly-GP staining is reduced in the nucleus of EDL muscle and D) Poly-GR staining is cleared from nuclei of EDL muscles after dox withdrawal. Single transgenic littermates, consisting of either TRE-only or rtTA-only, received the same dox treatment and are all negative for GFP and DPRs. All scale bars are 20 µm. All stainings were performed on all mice in this study. Numbers per group are: ST 1 week dox n=7, DT 1 week dox n=8, ST 1 week on/3 weeks off n=4, DT 1 week on/3 weeks off n=7, ST 2 weeks dox n=6, DT 2 weeks dox n=8, ST 2 weeks on/2 weeks off n=6, DT 2 weeks on/2 weeks off n=5, ST 4 weeks dox n=15, DT 4 weeks dox n=16.

**Figure 6:**
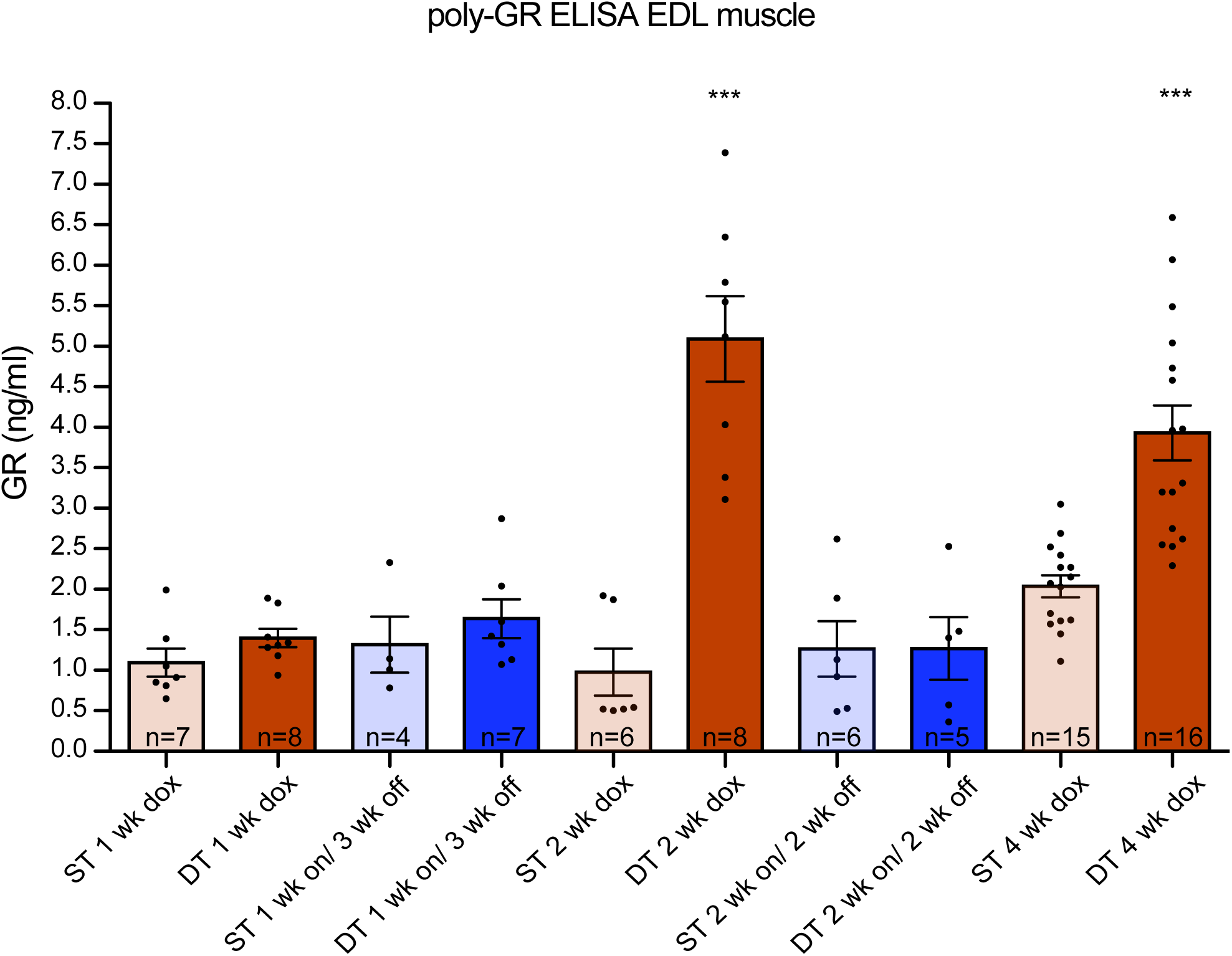
Expression of 36x G_4_C_2_human repeats *in vivo* generates high levels of poly-GR, which are reduced to single transgenic level after dox withdrawal. ELISA assessing poly-GR levels from mouse EDL muscle. Poly-GR is detectible in TRE-36xG4C2-GFP/hnRNP-rtTA double transgenic mice that received 2-4 weeks of dox. DT mice that received 1-2 weeks dox followed by 2-3 weeks washout have reduced amounts of poly-GR, similar to single transgenic levels. One-way ANOVA p=0.0001 Bonferroni post test shows that the DT 2 wk dox and DT 4 wk dox groups are significantly different than all the other groups, but not from each other. All other single transgenic and double transgenic washout groups are not significantly different from each other. Single transgenic littermates (consisting of only one transgene either TRE-only or rtTA-only) received similar dox treatment as double transgenic animals. Numbers per group are: ST 1 week dox n=7, DT 1 week dox n=8, ST 1 week on/3 weeks off n=4, DT 1 week on/3 weeks off n=7, ST 2 weeks dox n=6, DT 2 weeks dox n=8, ST 2 weeks on/2 weeks off n=6, DT 2 weeks on/2 weeks off n=5, ST 4 weeks dox n=15, DT 4 weeks dox n=16

### 36x G_4_C_2_ repeat mice develop a locomotor phenotype, rapid muscular dystrophy and NMJ abnormalities

Expression of 36x G_4_C_2_ repeats using the hnRNP-rtTA driver led to profound toxicity. We started with dox treatment in 6 weeks old mice to avoid DPRs affecting normal development, which would complicate behavioral and functional read-out. A large proportion (45%) of DT mice quickly declined in body weight in the first 2-3 weeks after dox administration and had to be sacrificed (figure 3A). Mice that quickly lost weight after 2.5 weeks showed general sickness symptoms (weight loss, bad condition of the fur, reduced activity, shivering) and an enlarged bladder. The majority of mice survived longer and did not lose weight but developed a locomotor phenotype on the Erasmus ladder (figure 3B and supplementary figure 8). This is a locomotor test based upon a horizontal ladder with alternating higher and lower rungs. Healthy C57Bl6/J mice prefer to walk on the higher rungs and avoid touching the lower rungs (Vinueza Veloz et al., 2015). DT mice began to touch the lower rungs more often after 2 weeks of dox treatment, although their performance was initially comparable to that of their ST littermates (figure 3B and supplementary figure 8) (two-way ANOVA analysis p=0.0001 for genotype and p<0.0001 for the interaction between genotype and time). DT mice sacrificed after 4 weeks of dox treatment displayed a white appearance of leg and back muscles macroscopically (figure 3C). At the histological level a massive distortion of muscle fibers could be observed (figure 3D). Histological analysis of other tissues revealed enlarged renal tubules in the kidney and hemorrhages in the bladder of mice that quickly lost weight at 2.5 weeks (figure 3D). Analyses of the neuromuscular junctions (NMJ) by whole mount immunostaining of the EDL muscle showed distortion of the muscular boutons and projecting motor neuron axons after 4 weeks of dox treatment (figure 3E). The number of motor neurons assessed by choline acetyltransferase (ChAT) staining of the spinal cord was not different between DT mice and ST control littermates (figure 3F). Together, these data indicate that body but no brain expression of 36x pure G_4_C_2_ repeats in our mouse model causes multi-system dysfunction, including urinary system problems and muscular dystrophy over the time course of one month.

### Early withdrawal of 36x G_4_C_2_ repeat expression can prevent but not reverse the muscular dystrophy phenotype

In order to investigate whether the phenotype could be reversed, we administered 6 weeks old DT and ST mice dox for 1 or 2 weeks and then changed them back to normal drinking water for 3 and 2 weeks, respectively (washout scheme figure 4A). Mice that only received 1 week of dox followed by 3 weeks of wash-out showed high survival rates and normal muscle and NMJ integrity (figure 4). About half of the DT mice that received 2 weeks of dox followed by 2 weeks of wash-out showed a fast reduction in body weight after 2-3 weeks and did not survive until the end of the experiment (figure 4B). This washout group was undistinguishable from the 4 weeks dox DT group regarding survival, muscle and NMJ integrity. Haematoxyline-eosine (HE) staining of the EDL muscle showed parts that displayed abnormal organization (figure 4C) and the NMJ showed disrupted boutons and axonal projections (figure 4D). Immunostaining for GFP, poly-GA and -GP in muscles of both washout groups showed clear reduction but was still detectable in nuclei of the 2 weeks on/ 2 weeks off group (figure 5). Only poly-GR could not be detected anymore (figure 5). In the kidney, GFP and all sense DPRs were cleared efficiently after dox withdrawal (supplementary figure 9). This indicates that DPR clearance is different per organ or cell type. To study build-up and reduction of DPRs in a more quantitative way, we developed an Enzyme-Linked Immuno Sorbent Assay (ELISA) for poly-GR, the DPR that shows the highest reported cellular toxicity. Poly-GR levels in EDL muscle increased significantly between the first and second week of dox administration, after which they stayed high (figure 6). In contrast, poly-GR levels in mice from the reversibility groups were reduced to similar levels as single transgenic control mice (one-way ANOVA test p=0.0001 with Bonferroni post test, figure 6). Together, our data shows that early withdrawal of 36x pure G_4_C_2_ repeat expression can prevent the accumulation of DPRs and the concurrent phenotype. Expression of 36x pure G_4_C_2_ repeat expression for 2 weeks and subsequent withdrawal for 2 weeks is not sufficient to completely clear DPRs and reverse muscular dystrophy *in vivo*.

## Discussion

In this study we demonstrate that expression of 36x pure G_4_C_2_ repeats *in vivo* is sufficient to cause NMJ abnormalities and muscular dystrophy leading to a specific locomotor phenotype within four weeks of transgene expression. Expression for four weeks did not evoke DPR expression in the brain, probably because four weeks are not long enough for build-up of DPRs and for doxycycline to pass the blood-brain barrier(Michel et al., 1984). Expression of 36x pure G_4_C_2_ repeats for 24 weeks in the murine brain, using a Camk2-alpha-rtTA driver, was also not sufficient to result in pathology or neurodegeneration, making this mouse model inadequate for studying brain specific DPR toxicity. We speculate that expression levels of the 36x G_4_C_2_ repeat RNA and DPRs in our Camk2-alpha-rtTA driven model are not high enough to induce neuropathology. Alternatively, the repeat length might not be long enough to evoke neurodegeneration. Other gain-of-function mouse models did show sense DPR pathology and a cognitive phenotype upon (over)expression of longer repeats(Chew et al., 2015; Herranz-Martin et al., 2017; Jiang et al., 2016; Liu et al., 2016). Mice expressing 500 repeats show a more severe phenotype compared with 29/36 repeat mice(Liu et al., 2016).

Upon expression of 36x pure G_4_C_2_ repeats in the body, our mouse model shows rapid muscular dystrophy and a locomotor phenotype. The phenotype could be caused by NMJ abnormalities or due to skeletal muscle dysfunction. Interestingly, DPR pathology has recently been found in skeletal muscle of C9ALS patients(Cykowski et al., 2019). DPR pathology has not been reported in tissues such as the kidney and bladder, even though C9ORF72 is expressed in these organs and C9KO mice show immune-mediated kidney damage (Atanasio et al., 2016; DeJesus-Hernandez et al., 2011). The pathology in our mouse model could be evoked by the relative rapid and strong repeat expression compared to the lower expression levels observed in C9FTD/ALS patients, but would be interesting to investigate how wide-spread DPR pathology is. Many *C9ORF72* mouse models lack locomotor symptoms due to unknown factors (Jiang et al., 2016; O’Rourke et al., 2015; Peters et al., 2015) and NMJ abnormalities have only been described in one BAC mouse model and one AAV-102x interrupted G_4_C_2_ mouse model (Herranz-Martin et al., 2017; Liu et al., 2016). Our mouse model shows similarities to the BAC 29/36 repeat mouse model reported by Liu et al. (Liu et al., 2016), as both models show DPR pathology but no RNA foci. However, the phenotype in our mouse model develops faster (within four weeks after dox administration) than reported by Liu et al. (first symptoms started after 16 weeks of age) (Liu et al., 2016). Differences in disease onset might be due to differences in expression levels. For example, lack of a phenotype was observed in a 37 repeat mouse with low expression levels(Liu et al., 2016), while our 36 repeat and the BAC 29/36 repeat mouse with higher expression levels clearly show a phenotype.

So far, the minimum repeat size to evoke RNA foci and DPR formation *in vivo* has remained unknown. A BAC mouse model of 110 repeats did not contain any RNA foci(Jiang et al., 2016), while BAC mice with longer repeat sizes did present with RNA foci (Jiang et al., 2016; Liu et al., 2016; O’Rourke et al., 2015; Peters et al., 2015). On the other hand, AAV-mediated overexpression of 10 or 66 repeats did evoke RNA foci(Chew et al., 2015; Herranz-Martin et al., 2017), indicating that formation of RNA foci could also depend on expression levels. Even though we did not detect any RNA foci in our 36x repeat mouse model, we cannot exclude an effect of repeat RNA on the observed phenotype. Repeat-containing RNA molecules might still be able to sequester molecules or proteins and affect their normal function of cellular processes.

Sense DPRs were detected as diffuse cytoplasmic or nuclear staining and did not form aggregates in our hnRNP-driven mouse model. Recent publications on poly-GR and -PR mouse models suggest that soluble poly-GR and -PR are sufficient to cause neurodegeneration and behavioral deficits(Zhang et al., 2018; Zhang et al., 2019). For poly-GA, aggregation seems necessary for its toxicity(Zhang et al., 2016). Thus, DPRs might differ in their abilities to aggregate, which can change their molecular targets and their effects on several cellular compartments and functions. Interestingly, poly-GA can spread throughout the brain and influence the aggregation of poly-GR and -PR(Darling et al., 2019; Moron-Oset et al., 2019; Yang et al., 2015), and this has been confirmed in AAV-66 and AAV-149x mice in which poly-GA and -GR co-aggregate in cells with poly-GA aggregates but poly-GR remains diffuse in cells devoid of poly-GA(Chew et al., 2019; Zhang et al., 2018). Poly-GA expression can even partially suppress poly-GR induced cell loss at the wing in a *Drosophila* model(Yang et al., 2015). Co-overexpression of poly-GA also abolished cellular toxicity of low concentrations of poly-PR in NSC34 cells(Darling et al., 2019). Other interactions between DPRs are still unknown and need further investigation. We did not detect antisense DPRs in our mouse model, maybe because antisense C_4_G_2_ RNA is not transcribed or antisense DPR levels might be too low to detect.

Another point of interest is the lack of apparent pTDP-43 pathology in multiple *C9ORF72* mouse models. TDP-43 pathology is thought to be a late event in the pathogenesis of C9FTD/ALS(Balendra and Isaacs, 2018). Several mouse models already show behavioral phenotypes and some mild neurodegeneration before the onset of pTDP-43 neuropathology(Herranz-Martin et al., 2017; Jiang et al., 2016; Schludi et al., 2017; Zhang et al., 2018; Zhang et al., 2016). Changes in pTDP-43 solubility or cellular localization could already arise and contribute to cellular distress without the formation of cytoplasmic aggregates per se(Lee et al., 2019). Indeed, several reports of C9FTD/ALS cases showed affected individuals with DPR pathology but mild or absent TDP-43 pathology(Baborie et al., 2015; Gijselinck et al., 2012; Mori et al., 2013; Proudfoot et al., 2014; Vatsavayai et al., 2016). Together, our hnRNP-driven mouse model shows that expression of diffuse labeled sense DPRs is sufficient to cause cellular toxicity without the need for RNA foci and pTDP-43 pathology.

The rapid translation of current knowledge into therapeutic intervention studies asks for robust *in vivo* drug discovery screens(Jiang and Ravits, 2019). So far, AON therapy has been tested in a BAC mouse model for *C9ORF72* and successfully reduced the amount of RNA foci and DPRs(Jiang et al., 2016). However, it remains unknown if this AON can also reduce motor symptoms associated with the *C9ORF72* repeat. New therapies are under development, including small molecules targeting RAN translation(Green et al., 2019; Hu et al., 2017; Kramer et al., 2016; Simone et al., 2018; Su et al., 2014; Zamiri et al., 2014; Zhang et al., 2015) and antibody therapy against poly-GA(Nguyen et al., 2020; Zhou et al., 2019). These therapies can be easily tested in our mouse model, as it develops a quick and robust phenotype. Our mouse model can be used as proof of principle for whole body toxicity of DPRs. We demonstrated that one week of expression followed by 3 weeks of washout (expression turned off) prevented the accumulation of DPRs and the associated cellular toxicity. However, two weeks of expression followed by two weeks of washout is not sufficient to prevent mice from developing muscular dystrophy. This indicates that transgene RNA or DPRs that were already produced during the first two weeks of dox administration continue to exercise their toxic effects. A recent publication estimated the half-lives of most DPRs to be >200 hours(Westergard et al., 2019). Half-life was longer for poly-GA puncta than for diffuse poly-GA and increased for poly-GR when localized in the nucleus(Westergard et al., 2019). Interestingly, poly-GP remains detectable after 1 week on/ 3 weeks off dox, indicating that poly-GP is not turning over as quickly as the other DPRs. Still, the animals of this reversibility group are improving, suggesting a bigger impact on toxicity of poly-GA and poly-GR, as shown before in earlier studies (Balendra and Isaacs, 2018; Mizielinska et al., 2014). In general, earlier intervention might be able to halt or reverse symptoms, but the preferred time-window for treatment is probably before the onset of symptoms.

Together, we provide evidence that expression of human 36x pure G_4_C_2_ repeats is sufficient to evoke RAN translation and a locomotor phenotype *in vivo*. Due to high expression of sense DPRs driven by hnRNP-rtTA, rapid progression of muscular dystrophy and NMJ disruption developed. This mouse model allows for fast *in vivo* screening of new drugs and compounds that act on systemic toxicity of sense DPRs.

## Material and Methods

### Cloning

DNA obtained from C9FTD patient 09D-5781 was assessed for the *C9ORF72* repeat expansion by Asuragen kit according to manufacturer’s protocol and contained at least 54 repeats. DNA was amplified in three consecutive rounds of PCR with primers flanking the *C9ORF72* repeat expansion (forward primer 5’-CCACGGAGGGATGTTCTTTA-3’ and reverse primer 5’-GAAACCAGACCCAAACACAGA-3’) and a PCR mix containing 50% betaine. The PCR program started with 10 min at 98 °C, followed by 35 cycles of 35 sec at 98 °C, 35 sec at 58°C and 3 minutes at 72°C and finished with 10 min at 72°C. The PCR product was cloned into TOPO vector PCR2.1 and restriction analysis with BsiEI (NEB) for 1 hour at 60 °C revealed a G_4_C_2_ repeat expansion estimated around 50 repeats. Next, the TRE-90CGG-GFP vector(Hukema et al., 2014) was restricted with SacII (NEB) and the 90xCGG repeat expansion was replaced with the 50x G_4_C_2_ repeat expansion. This vector was sequenced using a primer in the TRE sequence (5’-CGGGTCCAGTAGGCGTGTAC-3’) and revealed a repeat expansion of 36x G_4_C_2_. The final vector was cut with Aat II, PvuI and NdeI (NEB) and the band containing the TRE-36x G_4_C_2_-GFP construct was isolated from gel, dissolved in injection buffer (10 mM Tris-HCl, pH 7.4, 0.25 mM EDTA), and used to generate transgenic mice. Experiments on human material were done under informed consent and approved by the Medical Ethical Test Committee (METC).

### Animals

Pronuclei from oocytes of C57BL/6JRj wildtype (WT) mice were injected to create a new transgenic line harboring the TRE-36xG_4_C_2_-GFP construct. Genotyping was performed using primers located in the 5’ of the repeat expansion: forward 5’-GGTACCCGGGTCGAGGTAGG-3’ and reverse 5’-CTACAGGCTGCGGTTGTTTCC-3’. Founder mice, F1 and F2 were screened in an animal welfare assessment by the local animal caretakers and scored normal for litter size and health characteristics. All mice were housed in groups of 2 to 4 and were allowed to have free access to standard laboratory food and water. They were kept on a 12h light/dark cycle. TRE-36xG_4_C_2_-GFP mice were crossed with hnRNP-rtTA(Katsantoni et al., 2007) or Camk2-alpha-rtTA (kind gift of Rob Berman, Davis, USA) on a C57BL/6JRj WT background. Offspring should include 25% of double transgenic mice (harboring both the TRE- and one of the rtTA-constructs), 50% of single transgenic littermates (harboring either the TRE- or the rtTA-construct) and 25% of WT littermates (having no transgene). At 6 weeks of age, mice of both sexes were exposed to doxycycline (dox) (Sigma) (4 grams/L) combined with sucrose (50grams/L) dissolved in drinking water. Both single and double transgenic mice received dox water. To monitor the health and wellbeing, mice were weighed every weekday while on dox water (data not shown). TRE-36xG_4_C_2_-GFP/hnRNP-rtTA mice were sacrificed by cervical dislocation after maximal 4 weeks of dox administration. The TRE-36xG_4_C_2_-GFP/Camk2-alpha-rtTA mice were sacrificed by cervical dislocation after maximal 24 weeks of dox administration. As required by Dutch legislation, all experiments were approved in advance by the institutional Animal Welfare Committee (Erasmus MC, Rotterdam, The Netherlands).

### Erasmus Ladder

The Erasmus Ladder (Noldus, Wageningen, the Netherlands) is a fully automated test for detecting motor performance in mice(Vinueza Veloz et al., 2012). It consists of a horizontal ladder between two shelters, which are equipped with a bright white LED spotlight and pressurized air outlets. These are used as cues for departure from the shelter box to the other shelter box. The ladder has 2 x 37 rungs for the left and right side. All rungs have pressure sensors, which are continuously monitoring and registering the walking pattern of the mouse. The rungs are placed in an alternating high/low pattern. Wild type C57Bl6 mice prefer to walk on the higher rungs, avoiding touching the lower rungs [REF: Vinueza Veloz et al., Brain Structure and Function 2015]. The mouse was placed in the starting box and after a period varying from 9 to 11 seconds the LED light turned on and the mouse was supposed to leave the box. If the mouse left the box before the light turned on a strong air flow drove the mouse back into the box, and the waiting period restarted. If the mouse did not leave the box within 3 seconds after the light turned on, a strong air flow drove the mouse out of the box. When the mouse arrived in the other box the lights and air flow turned off and the waiting period from 9 to 11 seconds started and the cycle repeated again, making mice run back and forth on the ladder. Mice were trained on the Erasmus Ladder at the age of 5 weeks, every day for 5 days. The mice were trained to walk the ladder for 42 runs each day. At the age of 6 weeks the mice received dox/sucrose water and were tested on Monday, Wednesday and Friday on the Erasmus ladder. The average percentage of lower rung touches was calculated over 42 runs per session.

### Neuromuscular Junction staining

Extensor digitorum longus (EDL) muscles were fixed in 1% paraformaldehyde overnight (o/n). The muscles were washed in phosphate-buffered saline (PBS) and permeabilized in 2.5% Triton-X100 (Sigma) in PBS for 30 minutes and incubated in 1µg/ml α-bungarotoxin-TRITC (Invitrogen) in 1M NaCl for 30 minutes. Subsequently, muscles were incubated for 1 hour in a blocking solution (4% bovine serum albumin, 0.5% Triton-X100). After blocking, the muscles were incubated with a polyclonal chicken anti-neurofilament antibody (2Bscientific) 1:500 in blocking solution o/n at 4°C, followed by incubation for 4 hours with anti-chicken-alexa fluor 488 (Jackson Immuno Labs). Finally, the muscles were mounted on slides with 1.8% low-melting point agarose (ThermoFisher) and images were taken with a Zeiss LSM700 confocal microscope. The first author was blinded during image acquisition.

### Fluorescent In Situ Hybridization

Brain and EDL muscle tissues of mice were fixed in 4% paraformaldehyde o/n. Tissues were dehydrated and embedded in paraffin and cut into 6µm thick sections using a rotary microtome. Post-mortem human C9FTD/ALS frontal cortex paraffin tissue was used as positive control for RNA foci detection. Sections were deparaffinized using xylene and rehydrated in a standard alcohol series. Antigen retrieval was established in 0.01M sodium citrate with pH 6 using microwave treatment of 1x 9min followed by 2x 3min at 800W. Subsequently, the slides were dehydrated in an alcohol series and shortly dried to air. Next, pre-hybridization was performed in hybridization solution (dextran sulphate 10%w/v, formamide 50%, 2x SSC) for 1 hour at 65°C. After pre-hybridization, hexanucleotide sense oligo (5’-Cy5-4xGGGGCC-3’) and hexanucleotide antisense oligo (5’-Cy5-4xCCCCGG-3’) probes (IDT) were diluted to 40nM in hybridization solution and heated to 95°C for 5 minutes. The slides were hybridized with probe mix o/n at 65°C. After hybridization the slides were washed once with 2xSSC/0.1%Tween-20 and three times with 0.1x SSC at 65°C. Subsequently, slides were stained with Hoechst (Invitrogen), washed with PBS and stained with Sudan Black (Sigma). Finally, slides were dehydrated and mounted using Pro-Long Gold mounting solution (Invitrogen) and images were taken with a Zeiss LSM700 confocal microscope. C9FTD/ALS and non-demented control human brain sections were provided by the Dutch Brain Bank. Experiments on human material were done under informed consent and approved by the Medical Ethical Test Committee (METC).

### Immunohistochemistry

Tissues of mice were fixed in 4% paraformaldehyde o/n, and dehydrated and embedded in paraffin. 6µm thick sections were cut using a rotary microtome. Sections were deparaffinized using xylene and rehydrated in an alcohol series. Antigen retrieval was established in 0.01M sodium citrate with pH 6 using microwave treatment of 1x 9min followed by 2x 3min 800W. Endogenous peroxidase activity was blocked with 3% H_2_O_2_ and 1.25% sodium azide. Immunostaining was performed overnight at 4°C in PSB block buffer (1xPBS/0.5%protifar/0.15%glycine) and with the primary antibodies (see supplementary table 1 for all antibodies used in this study). The next day, sections were washed with PBS block buffer and antigen-antibody complexes were visualized by incubation with DAB substrate (DAKO) after incubation with Brightvision poly-HRP-linker (Immunologic) or anti-mouse/rabbit HRP (DAKO). Slides were counterstained with Mayer’s haematoxylin and mounted with Entellan (Merck Millipore International). The slides were imaged using a Olympus BX40 microscope (Olympus). C9FTD/ALS and non-demented control human brain sections were provided by the Dutch Brain Bank. Experiments on human material were done under informed consent and approved by the Medical Ethical Test Committee (METC).

### Protein isolation

Prior to lysing, EDL muscle samples were thawed on ice and supplied with RIPA buffer containing 0.05% protease inhibitors (Roche) and 0.3% 1M DTT (Invitrogen). Samples were mechanically lysed, followed by 30 min incubation on ice. After 30 min incubation, mechanical lysing was repeated and samples were centrifuged for 20 min at 4°C, followed by 3x 1 min sonication. After sonication, samples were centrifuged for 20 min at 4°C and the supernatant was used for ELISA. Whole protein content was determined using BCA assay (Thermo Fisher Scientific).

### Enzyme-Linked Immuno Sorbent Assay (ELISA)

MaxiSorp 96 well F-bottom plates (Thermo Fisher) were coated for 2h with 5.0 µg/ml monoclonal GR antibody followed by overnight blocking with 1% BSA in PBS-Tween (0.05% Tween-20, Sigma Aldrich) at 4°C. After washing, 300µg total protein lysate was added per sample. As positive control, a 15x GR synthetic peptide (LifeTein) was used. This peptide was serial diluted to create a standard curve (in duplo). All samples were measured both undiluted and diluted 2x and 4x with 0.1M PBS. Samples and GR peptide were incubated on the plate for 1h at RT. After washing, all wells were incubated for 1h with biotinylated monoclonal anti-GR antibody at a final concentration of 0.25 µg/ml in PBS-Tween/1% BSA. After washing again, samples were incubated for 20 min with Streptavidin-HRP (R&D Sciences) diluted 1:200 in PBS-Tween/1% BSA. Following extensive washing, samples were incubated with substrate reaction mix (R&D Sciences) for 15 min and stopped using 2N H2SO4. Read-out was carried out using a plate reader (Varioskan) at 450nm and 570nm.

### C9ORF72 Strand specific RT-PCR

RNA isolation was performed on mouse frozen kidney tissue and frozen frontal cortex of two C9FTD patients. Tissue was homogenized in 500 µl RIPA buffer (150 mM NaCl; 5mM EDTA; 50mM Tris-HCl, pH 8.0; 1% Nonidet-P40; 0.5 % sodium deoxycholate; 0.1% SDS, pH 7.6) containing Complete protease inhibitor (Roche), 3 mM DTT (Invitrogen) and 40 units RNAse OUT (Invitrogen). RNA was isolated using Trizol reagent (Invitrogen), according to manufacturer’s instructions. Reverse transcriptase was performed in 250 ng of RNA using SuperScriptIII cDNA synthesis kit (Invitrogen) according to manufacturer’s instructions. RNA was treated with DNase before cDNA synthesis. The following C9ORF72 strand specific primers were used to generate cDNA: LK-ASORF-R:5’-CGACTGGAGCACGAGGACACTGACGAGTGGGTGAGTGAGGAG-3’ (after repeat)orLK-ASORF-R2:5’-CGACTGGAGCACGAGGACACTGAGTAGCAAGCTCTGGAACTCAGGAGTCG-3’ (before repeat). For C9 FTD patient samples PCR was performed with LK specific primer: 5’-CGACTGGAGCACGAGGACACTGA-3’ and reverse primer: 5’-AGTCGCTAGAGGCGAAAGC-3’. For mouse samples PCR was performed with LK specific primer: 5’-CGACTGGAGCACGAGGACACTGA-3’ and reverse primer: 5’-CTCCTCACTCACCCACTCG-3’. PCR program: 4 min at 94°C followed by 35 cycles of 45s at 94°C, 45s at 58°C, 90s at 72°C, followed by 6 min at 72°C. C9FTD/ALS and non-demented control human brain frozen sections were provided by the Dutch Brain Bank. Experiments on human material were done under informed consent and approved by the Medical Ethical Test Committee (METC).

## Acknowledgements

Authors would like to thank Rob Berman for the Camk2-alpha-rtTA driver and Lies-Anne Severijnen for her guidance in IHC studies and anti-neurofilament antibody. We also thank Leonard Petrucelli for providing an aliquot of his anti-PA antibody and Elize Haasdijk for an aliquot of ChAT antibody.

## Author contributions statement

F.W.R. designed and performed experiments, interpreted data and wrote the manuscript. E.C.vdT. and R.F.M.V. performed experiments and analyzed data. A.M. and R.K.H. generated transgenic animals. L.W.J.B designed and interpreted behavioral assays. R.K.H and R.W. gave advice on experiments and interpreted data. R.K.H, L.W.J.B and R.W. reviewed and edited the manuscript.

