## Supplementary figures and legends for "Inducible expression of human *C9ORF72* 36x G_4_C_2_ hexanucleotide repeats is sufficient to cause RAN translation and rapid muscular atrophy in mice"

```
Mouse   17 ----GGA--------ATTCGCCCTTGtttttcccaccctctctCTcccCa
               |||        .||      ||||||||||||||  |||||||||||
Human  151 TTTAGGAGGTGTGTGTTT-----TTGTTTTTCCCACC--CTCTCTCCCCA

        55 ctacttgctcCCtcacagtactcgctgagggtgaacaagaaaagacctga
           |||||||||  |||||||||||||||||||||||||||||||||||||||
       194 CTACTTGCT--CTCACAGTACTCGCTGAGGGTGAACAAGAAAAGACCTGA

       105 taaagattaaccagaagaaaacaaggagggaaacaaccgcagcctgtagc
           ||||||||||||||||||||||||||||||||||||||||||||||||||
       242 TAAAGATTAACCAGAAGAAAACAAGGAGGGAAACAACCGCAGCCTGTAGC

       155 aagctctggaactcaggagtcgcgcgctaggggccggggccggggccggg
           |||||||||||||||||||||||||||||
       292 AAGCTCTGGAACTCAGGAGTCGCGCGCTA---------------------

       205 gccggggccggggccggggccggggccggggccggggccggggccggggc

       321 --------------------------------------------------

       255 cggggccggggccggggccggggccggggccggggccggggccggggccg

       321 --------------------------------------------------

       305 gggccggggccggggccggggccggggccggggccggggccggggccggg

       321 --------------------------------------------------

       355 gccggggccggggccggggccggggccggggccggggccggggccggggc
                                      |||||||||||||||||||||||
       321 ---------------------------GGGGCCGGGGCCGGGGCCGGGGC

       405 gtggtcggggcggcccgggggcgggcccggggcggggctgcggttgcgg
           |||||||||||||||||||||||||||||||||||||||||||||||||
       344 GTGGTCGGGGCGGCCCGGGGGCGGGCCCGGGGCGGGGCTGCGGTTGCGG

       455 tgcctgcgcccgcggcggaggcgcaggcggtggcgagtgggtgagtg
           |||||||||||||||||||||||||||||||||||||||||||||||
       394 TGCCTGCGCCCGCGGCGGAGGCGCAGGCGGTGGCGAGTGGGTGAGTG

       505 aggaggc--------ggcggg-------------------cccggga--
           |||||||        ||||||                   .||||||
       444 AGGAGGCGGCATCCTGGCGGTGGCTGTTTGGGGTTCGGCTGCCGGGAAG
```

**Supplementary figure 1: Alignment of DNA sequence of mouse transgene (lower case) and human C9ORF72 sequence (upper case/ capital letters) surrounding the repeat expansion.** NCBI Reference Sequence: NG_031977.1. Our mouse model contains 118 bp upstream and 115 bp downstream human flanking region around the G4C2 repeat expansion.

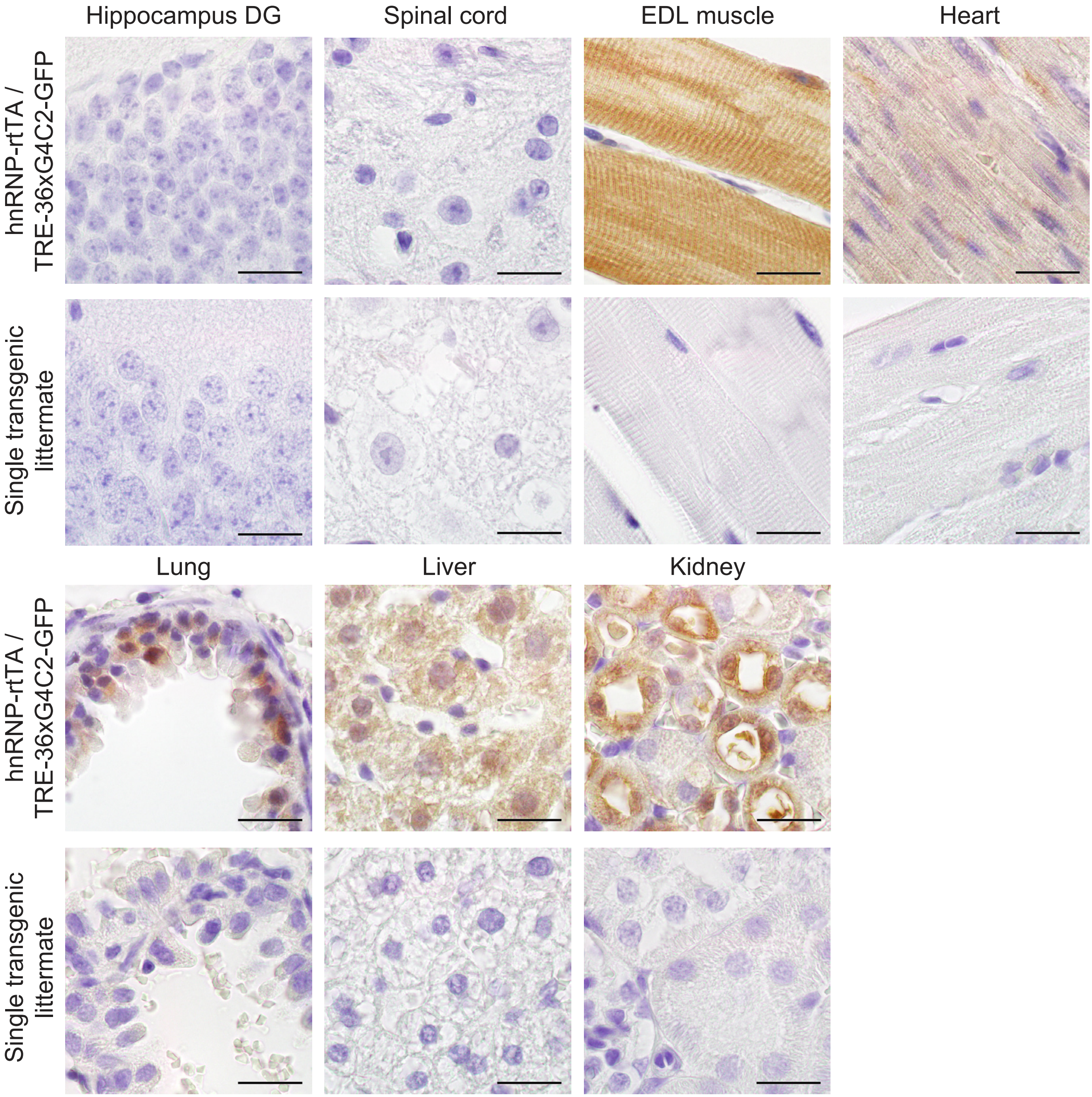


**Supplementary figure 2: GFP expression in EDL muscle, heart, lung, liver and kidney of TRE-36xG4C2-GFP/hnRNP-rtTA double transgenic mice.** No GFP staining was observed in the hippocampus dentate gyrus or in the spinal cord of DT mice. Single transgenic littermates, consisting of either TRE-only or rtTA-only, received the same dox treatment and are all negative for GFP staining. Scale bars are 20 µm. All stainings were performed on all mice in this study. ST 4 weeks dox n=15, DT 4 weeks dox n=16

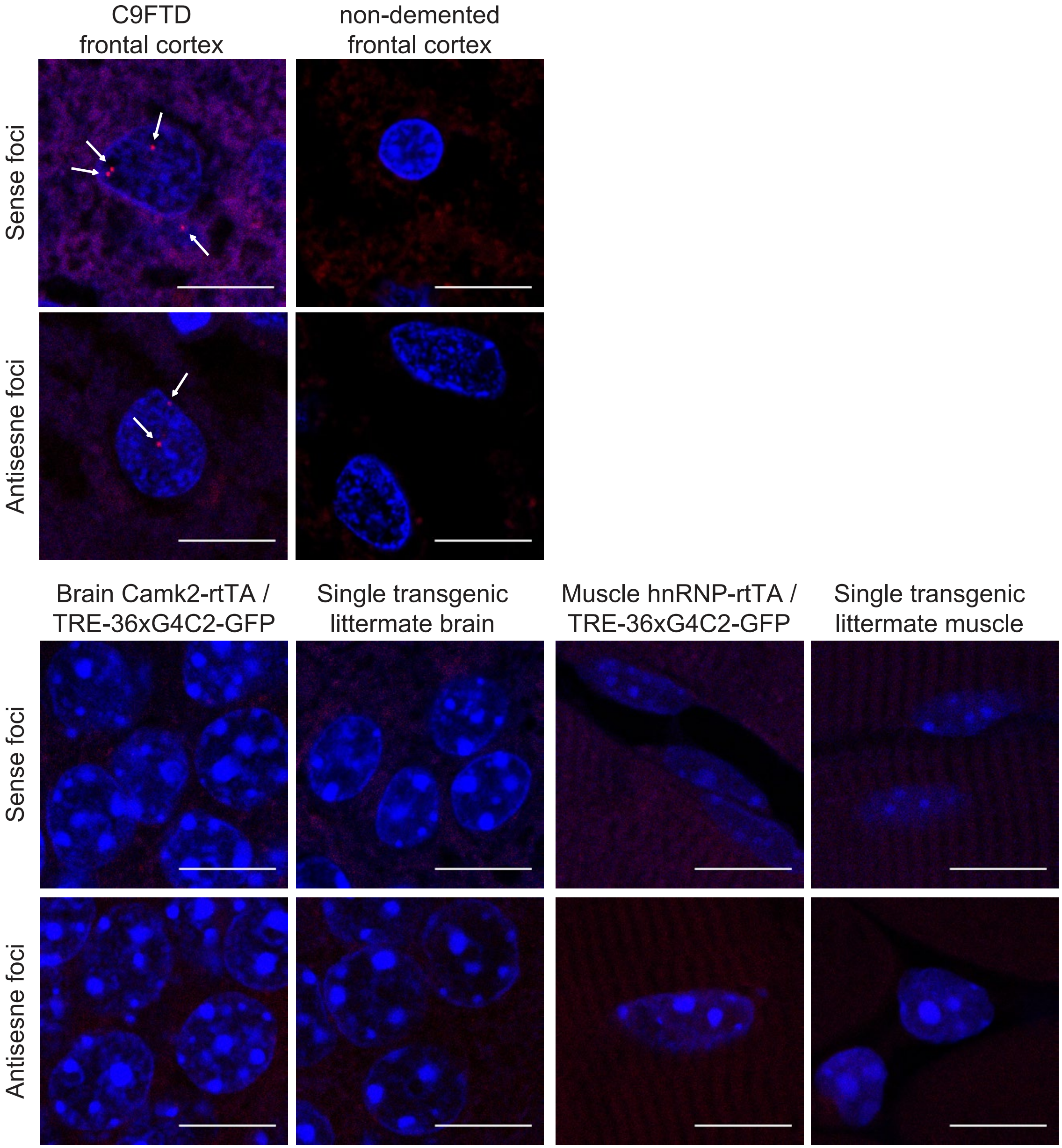


**Supplementary figure 3: No sense nor antisense RNA foci were found in TRE-36xG4C2-GFP/Camk2-alpha-rtTA and TRE-36xG4C2-GFP/hnRNP-rtTA double transgenic mice and control single transgenic littermates.** Single and double transgenic mice received the same dox treatment. Only frontal cortex samples of C9FTD cases present with some nuclear sense and antisense foci. Scale bars are 10 µm. The FISH was performed on all mice in this study. ST 4 weeks dox n=15, DT 4 weeks dox n=16

**A**

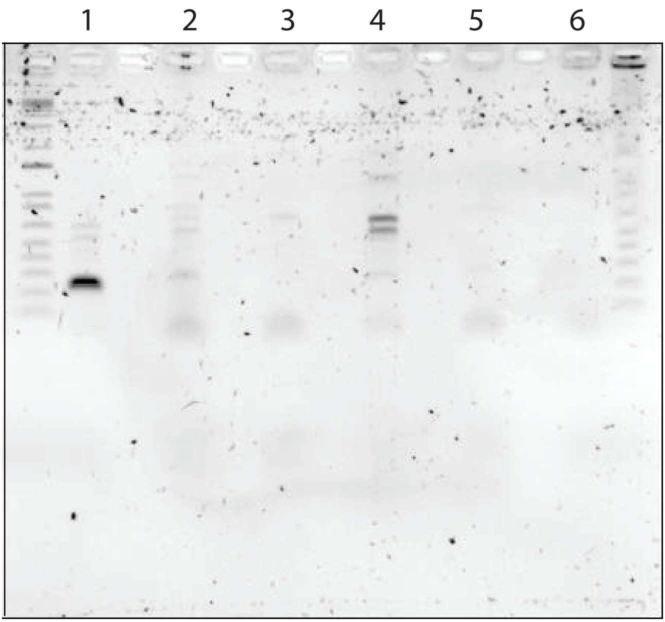


**B**

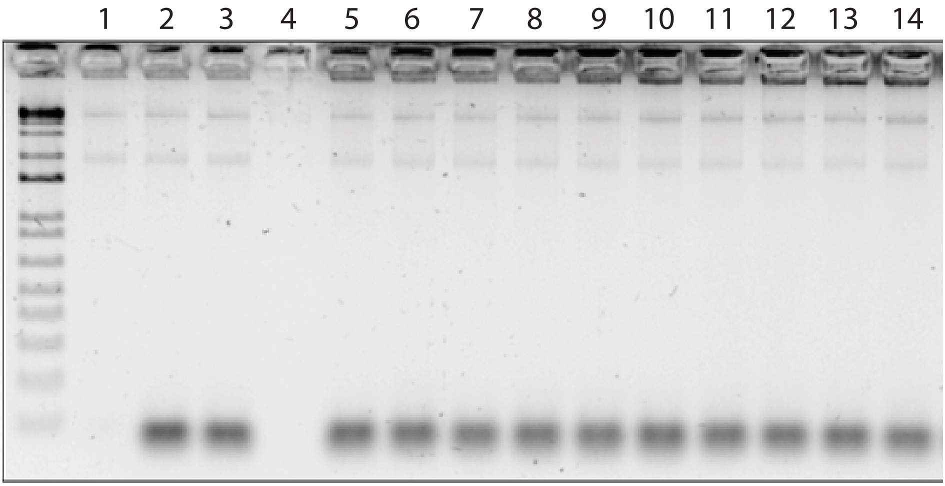


**Supplementary figure 4: reverse transcriptase PCR for C9orf72 antisense transcripts.** A) rt-PCR for C9orf72 antisense transcripts on frozen human prefrontal cortex samples with C9orf72 reverse specific primers. 1: NHB 05-123 (C9FTD) 2: NHB 09-044 (C9FTD) 3: NHB 08-150 (non-demented control) 4: NHB 07-089 (non-demented control) 5: NHB 05-123 (C9FTD) -rt control 6: H2O control. B) rt-PCR for C9orf72 antisense transcripts on TRE-36xG4C2-GFP mouse kidney samples with C9orf72 reverse specific primers. 1: H2O control 2: -rt mouse (15547-03 –rt) 3: -rt human (05-123 –rt) 4: empty cause of broken well 5: 15547-04 (DT) 6: 15547-03 (DT) 7: 12938-08 (DT) 8: 12938-06 (DT) 9: 18379-04 (DT) 10: 18379-03 (ST) 11: 18154-02 (DT) 12: 18154-01 (DT) 13: 18379-07 (DT) 14: 18379-01 (DT). Single and double transgenic mice received the same dox treatment.

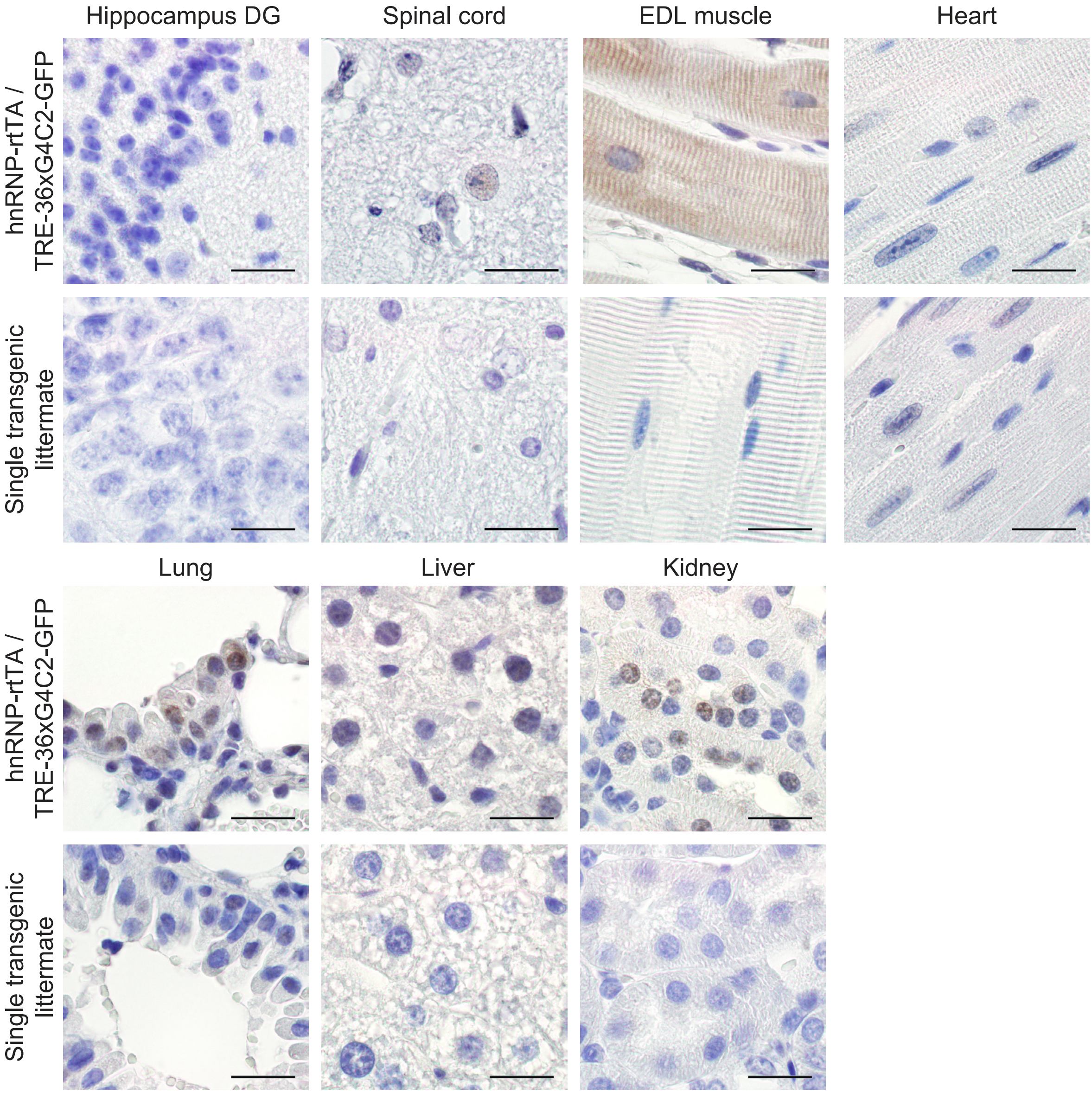


**Supplementary figure 5: Poly-GA expression in EDL muscle, lung, liver and kidney of TRE-36xG4C2-GFP/hnRNP-rtTA double transgenic mice.** No poly-GA staining was observed in the hippocampus dentate gyrus or in the spinal cord of DT mice. Single transgenic littermates, consisting of either TRE-only or rtTA-only, were treated similarly with dox and are all negative for poly-GA staining. Scale bars are 20 µm. The poly-GA was performed on all mice in this study. ST 4 weeks dox n=15, DT 4 weeks dox n=16

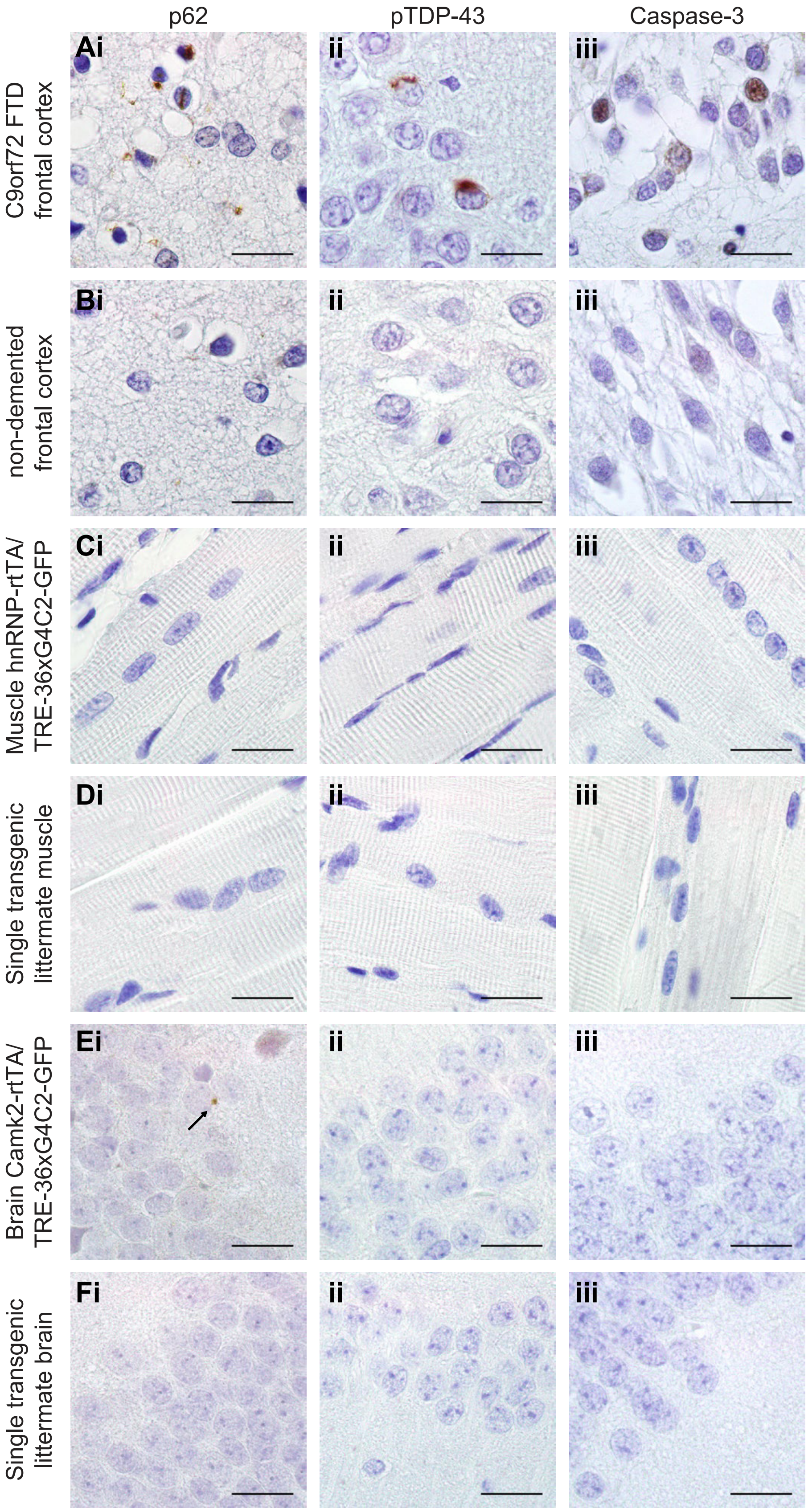


**Supplementary figure 6: Expression of 36x G4C2 human repeats does not cause abundant p62, pTDP-43 and cleaved-caspase 3 pathology.** Human prefrontal cortex of A) C9FTD patients or B) non-demented controls were used as positive and negative control for detection of pathology. C) TRE-36xG4C2-GFP/hnRNP-rtTA double transgenic mice do not present with any p62, pTDP-43 or cleaved-caspase-3 pathology in EDL muscle. E) TRE-36xG4C2-GFP/Camk2-alpha-rtTA double transgenic mice show some sparse perinuclear aggregates of p62 in the hippocampus dentate gyrus (arrow). D) and F) Single transgenic littermates, consisting of either TRE-only or rtTA-only, were treated similarly with dox and are negative for all pathology. All scale bars are 20 μm. All stainings were performed on all mice in this study. ST 4 weeks dox n=15, DT 4 weeks dox n=16.

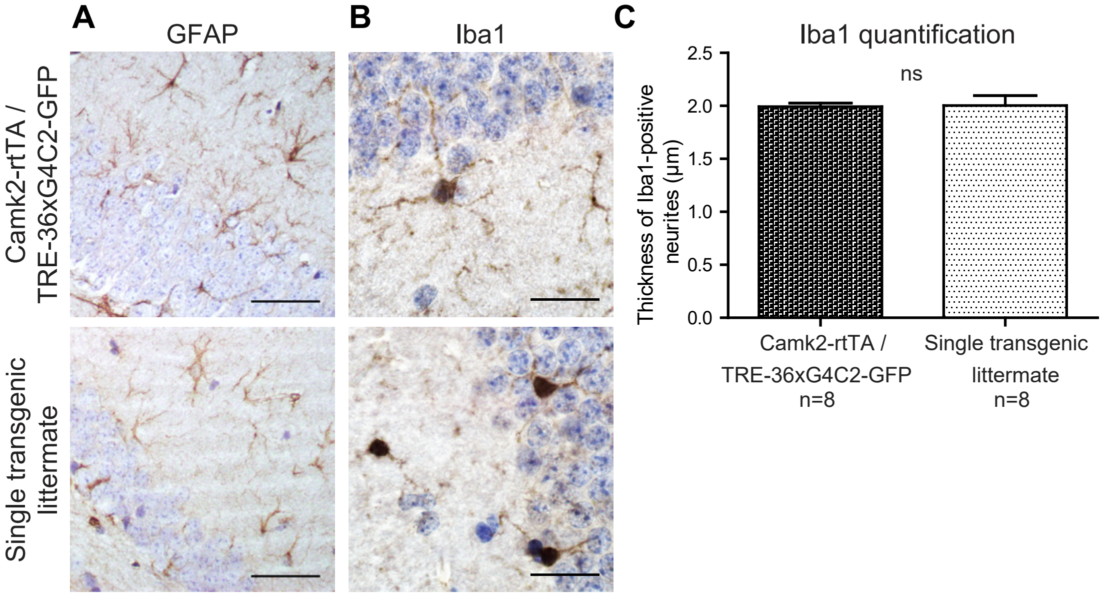


**Supplementary figure 7: TRE-36xG4C2-GFP/Camk2-alpha-rtTA double transgenic mice do not show astrogliosis or microgliosis.** A) Astrogliosis was assessed with GFAP labeling and B) microgliosis was tested with Iba1 staining. No differences in amount or morphology of GFAP-positive and Iba1-positive cells were seen in the hippocampus dentate gyrus of TRE-36xG4C2-GFP/Camk2-alpha-rtTA double transgenic mice and single transgenic control littermates. Single transgenic littermates, consisting of either TRE-only or rtTA-only, were treated similarly with dox. Scale bars are 20 µm. C) To quantify the thickness of Iba1-positive neurites, averages were taken of 10 pictures per mouse. N=8 TRE-36xG4C2-GFP/Camk2-alpha-rtTA double transgenic mice and n=8 single transgenic controls. T-test p=0.9099.

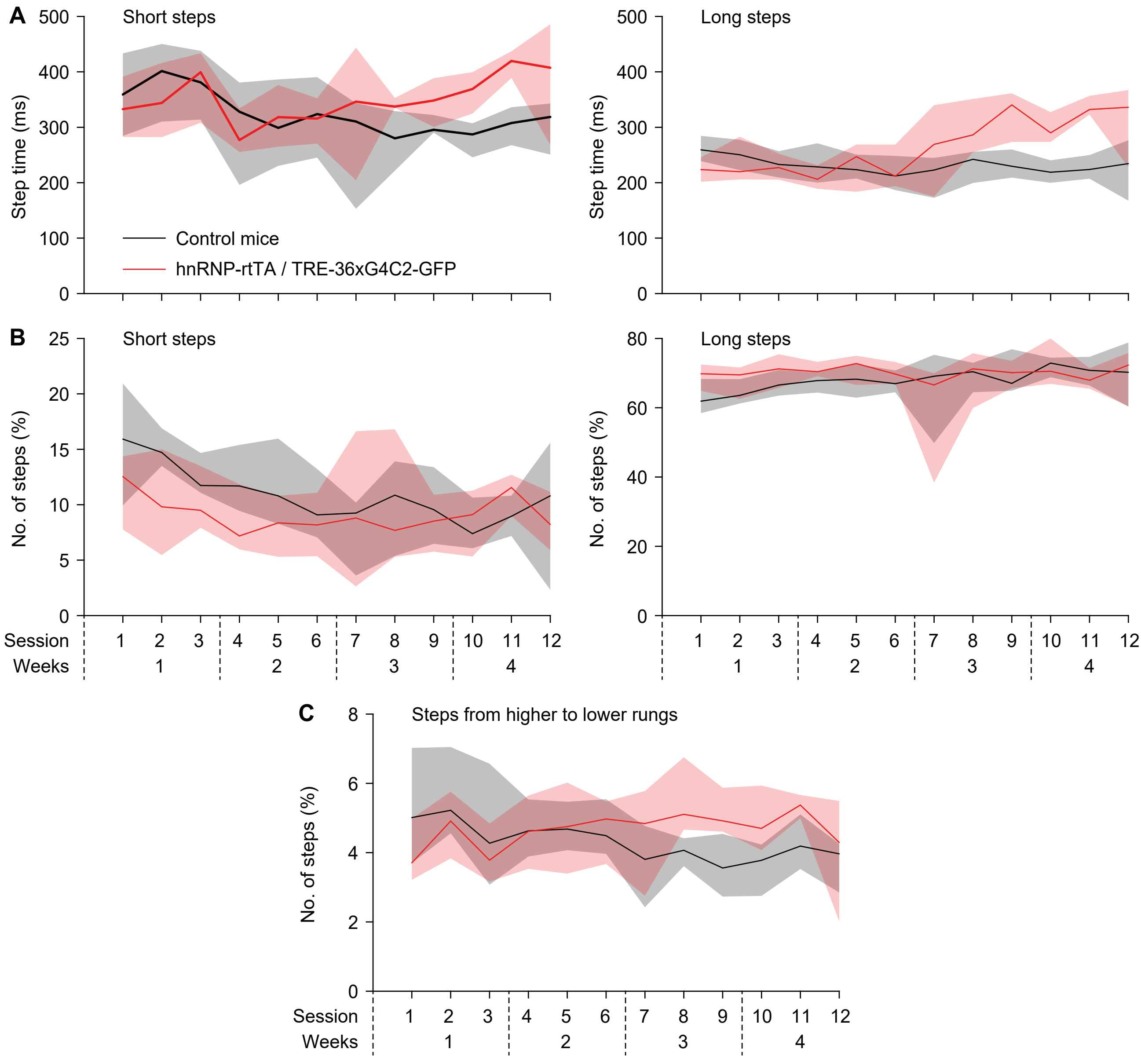


**Supplementary figure 8: Erasmus ladder readouts.** A) Step times of short steps (from one higher rung to the next; left) and long steps (skipping one higher rung, right). B) Fraction of short and long steps of all steps. C) Fraction of steps that were made from a higher rung to a lower rung. Lines indicate medians and shaded areas the interquartile ranges.

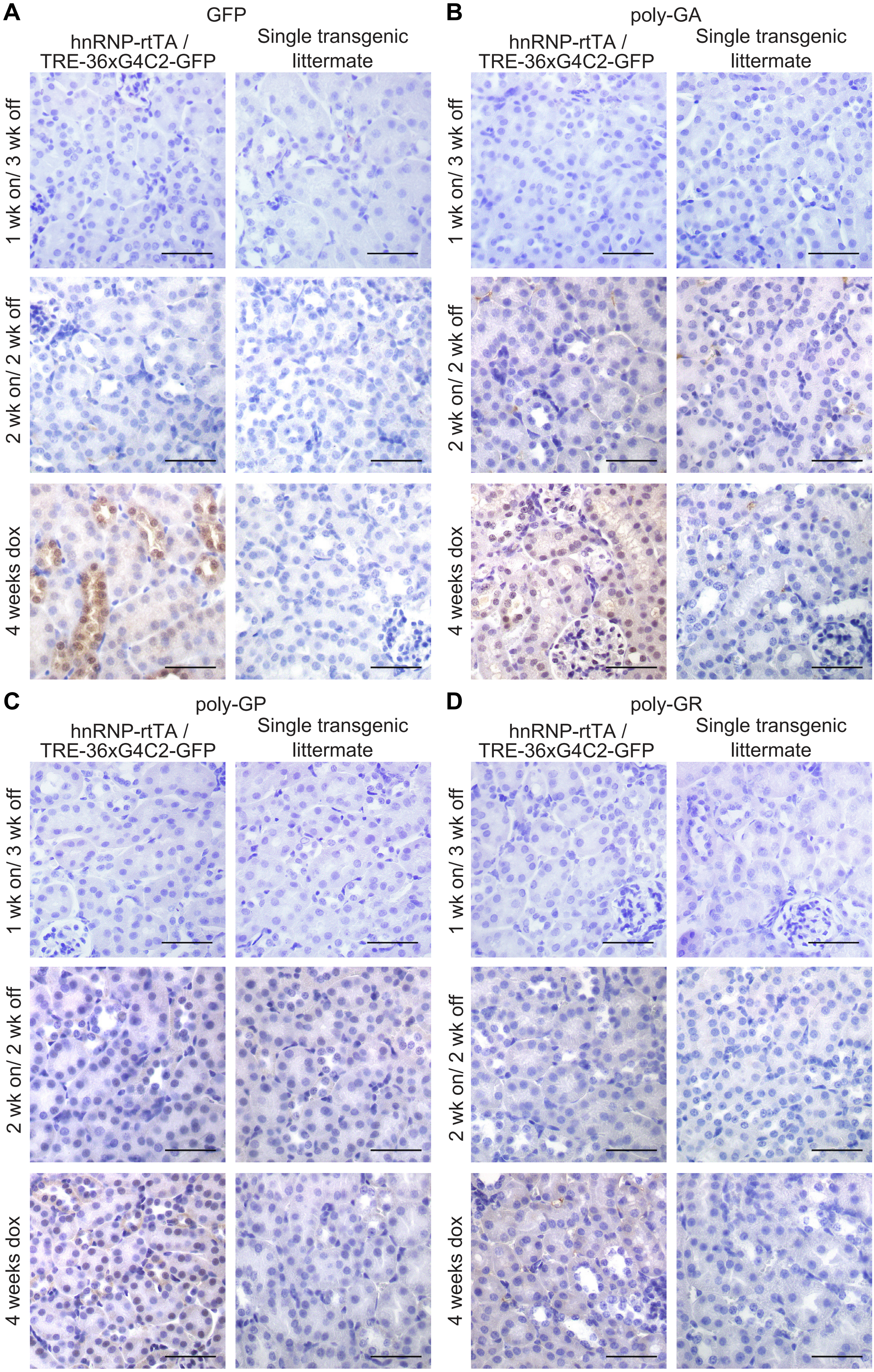


**Supplementary figure 9: GFP and sense DPRs are cleared from the kidney after 2 weeks of dox withdrawal.** A) GFP staining on kidney of TRE-36xG4C2-GFP/hnRNP-rtTA double transgenic mice shows clearance of GFP staining when mice received 2 weeks of dox water followed by 2 weeks of normal drinking water compared to DT littermates that received 4 weeks of dox. B) Poly-GA staining of kidney shows clearance of poly-GA after two weeks of dox withdrawal. C) Poly-GP staining and D) Poly-GR staining are also cleared from kidneys after 2 weeks of dox withdrawal. Single transgenic littermates received 2 or 4 weeks of dox and are all negative for GFP and DPRs. All scale bars are 50 µm. All stainings were performed on all mice in this study. Numbers per group are: ST 1 week dox n=7, DT 1 week dox n=8, ST 1 week on/3 weeks off n=4, DT 1 week on/3 weeks off n=7, ST 2 weeks dox n=6, DT 2 weeks dox n=8, ST 2 weeks on/2 weeks off n=6, DT 2 weeks on/2 weeks off n=5, ST 4 weeks dox n=15, DT 4 weeks dox n=16

**Supplementary table 1: antibodies**

| Ab name | Host | Company | Cat.nr | Dilution |
|---|---|---|---|---|
| GA | mouse | Millipore, clone 5E9 | MABN889 | 1:500 |
| GP | rabbit | Bio Connect Life Sciences | 24494-1-AP | 1:250 |
| GR | mouse | LifeTein Services | n.a. (costum-made) | 1:4000 |
| PR | mouse | LifeTein Services | n.a. (costum-made) | 1:500 |
| PA | mouse | Gift from Petrucelli | n.a. | 1:2500 |
| pTDP-43 | mouse | Cosmo bio | CAC-TIP-PTD-M01 | 1:1000 |
| p62 | mouse | BD Biosciences | 610833 | 1:100 |
| Neurofilament | chicken | 2BScientific Ltd. | CPCA-NF-H-25ul | 1:500 |
| GFAP | Rabbit | Sigma | G-9269 | 1:100 |
| Iba1 | rabbit | Wako | 019-19741 | 1:200 |
| ChAT | goat | Chemicon | AB144P | 1:500 |
| poly-HRP anti Ms/Rb IgG | goat | Immunologic | DPV055HRP | undiluted |
| anti-mouse HRP | goat | DAKO | P0260 | 1:100 |
| anti-rabbit HRP | goat | DAKO | P0217 | 1:100 |
| anti-mouse Cy2 | goat | Jackson | 715-255-150 | 1:100 |
| anti-rabbit Cy3 | goat | Jackson | 711-165-152 | 1:100 |
| anti-chicken 488 | goat | Jackson | 303-545-006 | 1:100 |
| anti-goat HRP | rabbit | DAKO | P0449 | 1:100 |
